# Ultrasonographic measurements of fascicle length overestimate adaptations in serial sarcomere number

**DOI:** 10.1101/2023.06.02.543410

**Authors:** Avery Hinks, Martino Franchi, Geoffrey A. Power

## Abstract

Ultrasound-derived measurements of muscle fascicle length (FL) are often used to infer increases (chronic stretch or training) or decreases (muscle disuse or aging) in serial sarcomere number (SSN). Whether FL adaptations measured via ultrasound can truly approximate SSN adaptations has not been investigated. We casted the right hindlimb of 15 male Sprague-Dawley rats in a dorsiflexed position (i.e., stretched the plantar flexors) for 2 weeks, with the left hindlimb serving as a control. Ultrasound images of the soleus, lateral gastrocnemius (LG), and medial gastrocnemius (MG) were obtained with the ankle at 90° and full dorsiflexion for both hindlimbs pre and post-cast. Following post-cast ultrasound measurements, legs were fixed in formalin with the ankle at 90°, then muscles were dissected, and fascicles were teased out for measurement of sarcomere lengths via laser diffraction and calculation of SSN. Ultrasound detected an 11% increase in soleus FL, a 12% decrease in LG FL, and an 8-11% increase in MG FL for proximal fascicles and at full dorsiflexion. These adaptations were partly reflected by SSN adaptations, with a 6% greater soleus SSN in the casted leg than the un-casted leg, but no SSN differences for the gastrocnemii. Weak relationships were observed between ultrasonographic measurements of FL and measurements of FL and SSN from dissected fascicles. Our results showed that ultrasound-derived FL measurements can overestimate an increase in SSN by ∼5%. Future studies should be cautious when concluding a large magnitude of sarcomerogenesis from ultrasound-derived FL measurements, and may consider applying a correction factor.

**Key Points Summary:**

- Measurements of muscle fascicle length via ultrasound are often used to infer changes in serial sarcomere number, such as increases following chronic stretch or resistance training, and decreases with aging or muscle disuse
- The present study used a rat model of casting the plantar flexor muscles in a stretched position to investigate directly whether ultrasound-derived fascicle length can accurately detect adaptations in serial sarcomere number
- Ultrasound detected an ∼11% increase in soleus fascicle length, but measurements on dissected fascicles showed the actual increase in serial sarcomere number was only ∼6%; therefore, measurements of ultrasound-derived fascicle length can overestimate serial sarcomere number adaptations by as much as 5%

## Introduction

Characterization of a muscle’s serial sarcomere number (SSN) gives insight into properties of biomechanical function (Lieber & Fridén, 2000; Narici *et al*., 2016; Hinks *et al*., 2022*a*). To that end, B-mode ultrasound is often used in humans to measure fascicle length (FL) and infer SSN adaptations at a smaller scale, such as increases in FL following resistance training (Blazevich *et al*., 2007; Franchi *et al*., 2014; Hinks *et al*., 2021) or decreases in FL with age and disuse (Williams & Goldspink, 1978; Narici *et al*., 2003; de Boer *et al*., 2008; Power *et al*., 2013). In animals, SSN can be estimated more precisely by dividing average sarcomere length (SL) measured via laser diffraction by the length of a dissected fascicle (Butterfield *et al*., 2005; Chen *et al*., 2020; Hinks *et al*., 2022*b*). Unfortunately, direct measurement of SL in humans is invasive (Lieber *et al*., 1997; Boakes *et al*., 2007), and often prohibitively costly and not accessible (Lichtwark *et al*., 2018; Adkins *et al*., 2021). However, inferring SSN adaptations via ultrasound-derived measurements of FL may be problematic because apparent increases or decreases in FL could be due to longer or shorter SLs, respectively, at the joint angle in which FL was measured (Pincheira *et al*., 2021). The relationship between SSN and FL may also depend on the region of muscle, with the human tibialis anterior displaying greater SSN in proximal fascicles due to a shorter SL (Lichtwark *et al*., 2018). Collectively, the relationship between SSN and ultrasound-derived FL may depend on the joint angle and region of muscle at which measurements are taken. Whether FL adaptations measured via ultrasound truly approximate SSN adaptations has not been investigated.

Assessment of FL in rodents via ultrasound is less common than in humans, but not unfounded. Peixinho and colleagues developed reliable methods for assessment of muscle architecture via ultrasound in the rat plantar flexors (Peixinho *et al*., 2011, 2014). Ultrasonography of the rat plantar flexors also has enough sensitivity to detect morphological adaptations (Peixinho *et al*., 2014; Mele *et al*., 2016). These previous studies, however, only assessed pennation angle (PA) and muscle thickness, leaving characterization of ultrasound-derived FL adaptations in rats unclear. Altogether, rodent models present an opportunity to assess the sensitivity of ultrasound measurements of FL in detecting actual SSN adaptations.

The present study assessed the validity of ultrasound as a tool to detect adaptations in SSN. To do this, we immobilized the rat plantar flexors in a lengthened position—an intervention that rapidly increases soleus SSN (Tabary *et al*., 1972; Williams & Goldspink, 1978; Soares *et al*., 2007; Aoki *et al*., 2009). We hypothesized that the ability for ultrasound-derived FL measurements to characterize adaptations in SSN would vary depending on the joint angle at which ultrasound measurements are obtained and the region of muscle.

## Methods

### Animals

Fifteen male Sprague-Dawley rats (sacrificial age ∼19 weeks) were obtained (Charles River Laboratories, Senneville, QC, Canada). Approval was given by the University of Guelph’s Animal Care Committee and all protocols followed CCAC guidelines (AUP #4905). Rats were housed at 23°C in groups of three and given ad-libitum access to a Teklad global 18% protein rodent diet (Envigo, Huntington, Cambs., UK) and room-temperature water. The right leg was immobilized in dorsiflexion for 2 weeks to place the plantar flexor muscles, in particular the soleus, in a lengthened position (Soares *et al*., 2007; Aoki *et al*., 2009). Per previous investigations of SSN adaptations in immobilized rat muscle, the contralateral limb served as a control (Heslinga & Huijing, 1993; Gomes *et al*., 2004). Ultrasound images of the lateral gastrocnemius (LG), medial gastrocnemius (MG), and soleus were obtained at ∼17 weeks of age (pre-immobilization) and ∼19 weeks of age (post-immobilization).

### Unilateral Immobilization

Using gauze padding, vet wrap, and a 3D-printed brace and splint, the right hindlimb of each rat was immobilized in dorsiflexion (40° ankle angle; full plantar flexion = 180°) (Figure 1A). Casts were inspected daily and repaired/replaced as needed. The toes were left exposed to monitor for swelling (Aoki *et al*., 2009).

**Figure 1:**
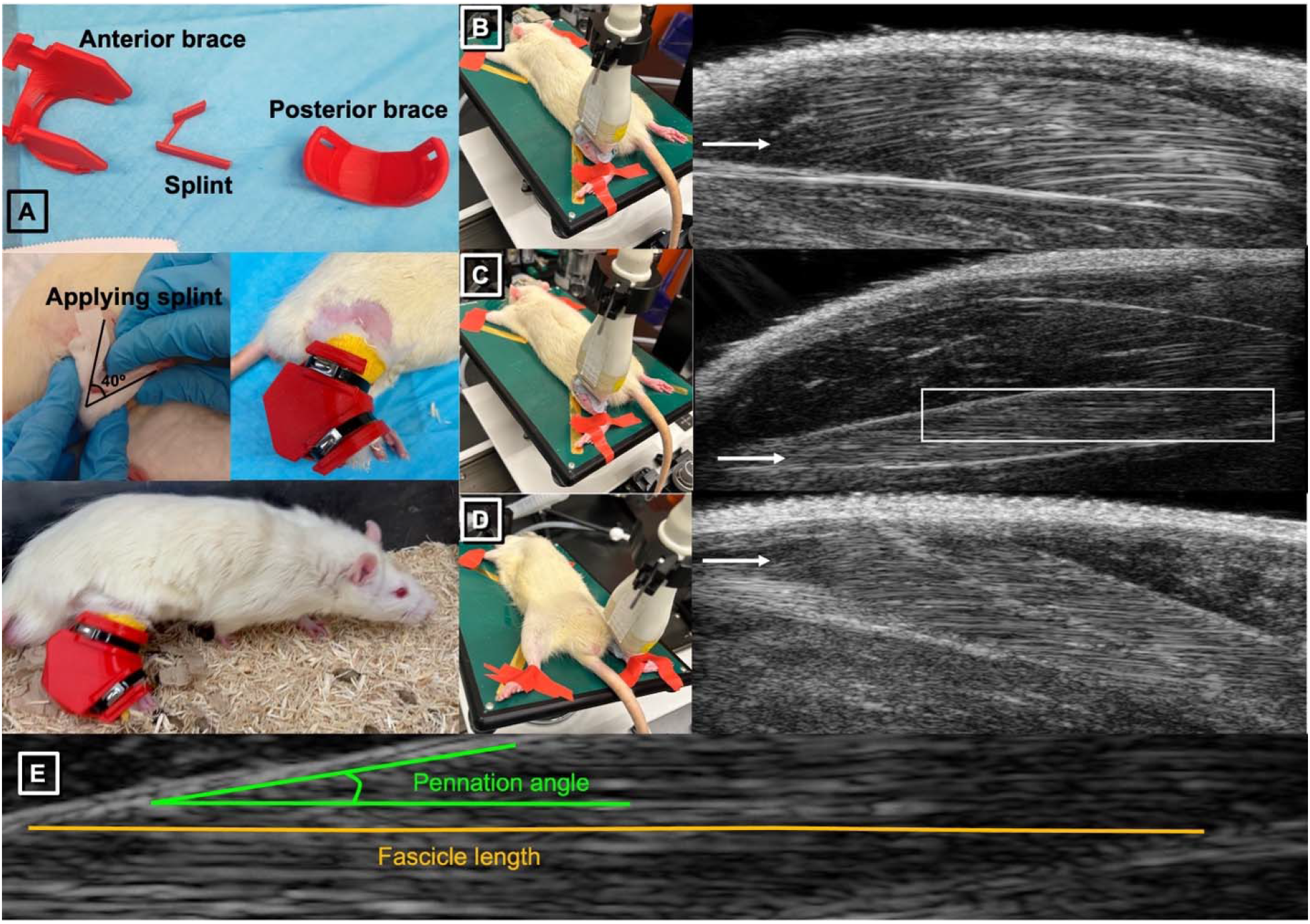
**A.** Example images of applying the splint and brace for the dorsiflexion cast. **B-D.** Setup and example of ultrasound images obtained from the left lateral gastrocnemius (B), soleus (C), and medial gastrocnemius (D), with the ankle fixed at 90° using tape. White arrows indicate the muscle of interest in each image. **E** corresponds to the area highlighted by the white box in C and shows representative tracings of fascicle length (orange) and pennation angle (green).

### Ultrasonography

Ultrasound measurements were obtained from the right and left hindlimbs at pre-immobilization (no more than 1 week prior to first applying the casts) and post-immobilization (immediately following cast removal).

A UBM system (Vevo 2100; VisualSonics, Toronto, ON, Canada) operating at a centre frequency of 21 MHz was used to acquire images of the soleus, LG, and MG, with a lateral resolution of 80 μm and an axial resolution of 40 μm (Mele *et al*., 2016). A 23-mm long probe was used, allowing acquisition of images displaying muscle fascicles from end to end. During piloting, image acquisition was optimized with an image depth of 15 mm for the soleus and LG and 16 mm for the MG, both allowing a maximum frame rate of 16 Hz. Prior to image acquisition, rats were anesthetized using isoflurane. With the knee fully extended, tape was used to fix the ankles at two different positions for image acquisition: 1) 90°; and 2) full dorsiflexion. All ultrasound images were acquired by the same individual (A.H.). Images of the LG and soleus were obtained with the rat in a prone position and the hindlimb externally rotated, with the probe overlying the lateral aspect of the posterior shank (Figure 1B-C). Images of the MG were obtained with the rat in a supine position and the hindlimb externally rotated, with the probe overlying the medial aspect of the posterior shank (Figure 1D). The probe position was carefully adjusted to obtain the clearest possible view of fascicles in all of the proximal, middle, and distal regions of the muscle. Throughout image acquisition, the probe was stabilized by a crane with fine-tune adjustment knobs, minimizing pressure and limiting the error associated with human movement.

Ultrasound images were analysed using ImageJ software (Franchi *et al*., 2020). ImageJ’s multisegmented tool allowed careful tracing of the fascicle paths from end to end in measuring FL. Two measurements of FL and PA were obtained from each of the proximal, middle, and distal regions of each muscle (i.e., six FL and PA measurements per muscle). PA was defined as the angle between the fascicle and the aponeurosis at the fascicle’s distal insertion point. All FL and PA measurements were obtained by the same experimenter (A.H.), who was blinded to the results until all measurements pre-and post-immobilization were obtained. During piloting, across three separate image acquisitions on the same rat, the coefficients of variation (standard deviation / mean × 100%) for FL averaged among two measurements at each region of muscle were all <10% (Table 1), which indicates low variation among repeated measures.

**Table 1:**
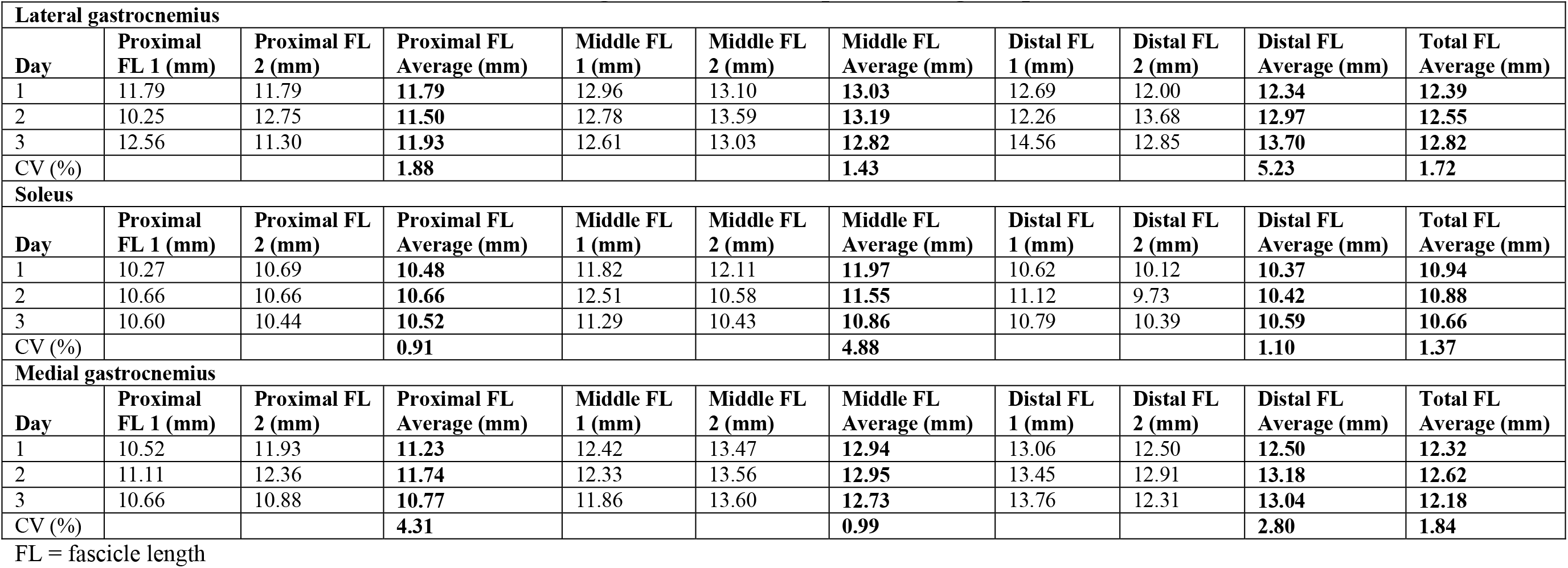
Coefficients of variation for fascicle length across three separate image acquisitions on the same rat.

### Serial Sarcomere Number Estimations

Following the post-immobilization ultrasound measurements, rats were sacrificed, and the hindlimbs were amputated and fixed in 10% phosphate-buffered formalin with the ankle pinned at 90° and the knee fully extended. After fixation for 1-2 weeks, the muscles were dissected and rinsed with phosphate-buffered saline. The muscles were then digested in 30% nitric acid for 6-8 hours to remove connective tissue and allow for individual muscle fascicles to be teased out (Butterfield *et al*., 2005; Hinks *et al*., 2022*b*).

For each muscle, two fascicles were obtained from each of the proximal, middle, and distal regions of the muscle (i.e., six fascicles total per muscle). Dissected fascicles were placed on glass microslides (VWR International, USA), then FLs were measured using ImageJ software (version 1.53f, National Institutes of Health, USA) from pictures captured by a level, tripod-mounted digital camera, with measurements calibrated to a ruler in plane with the fascicles (Supplemental Figure S1). Sarcomere length measurements were taken at six different locations proximal to distal along each fascicle via laser diffraction (Coherent, Santa Clara, CA, USA) with a 5-mW diode laser (25 μm beam diameter, 635 nm wavelength) and custom LabVIEW program (Version 2011, National Instruments, Austin, TX, USA) (Lieber *et al*., 1984), for a total of 36 sarcomere length measurements per muscle. Serial sarcomere numbers was calculated as:

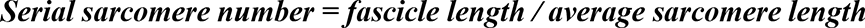

**Supplemental Figure S1:**
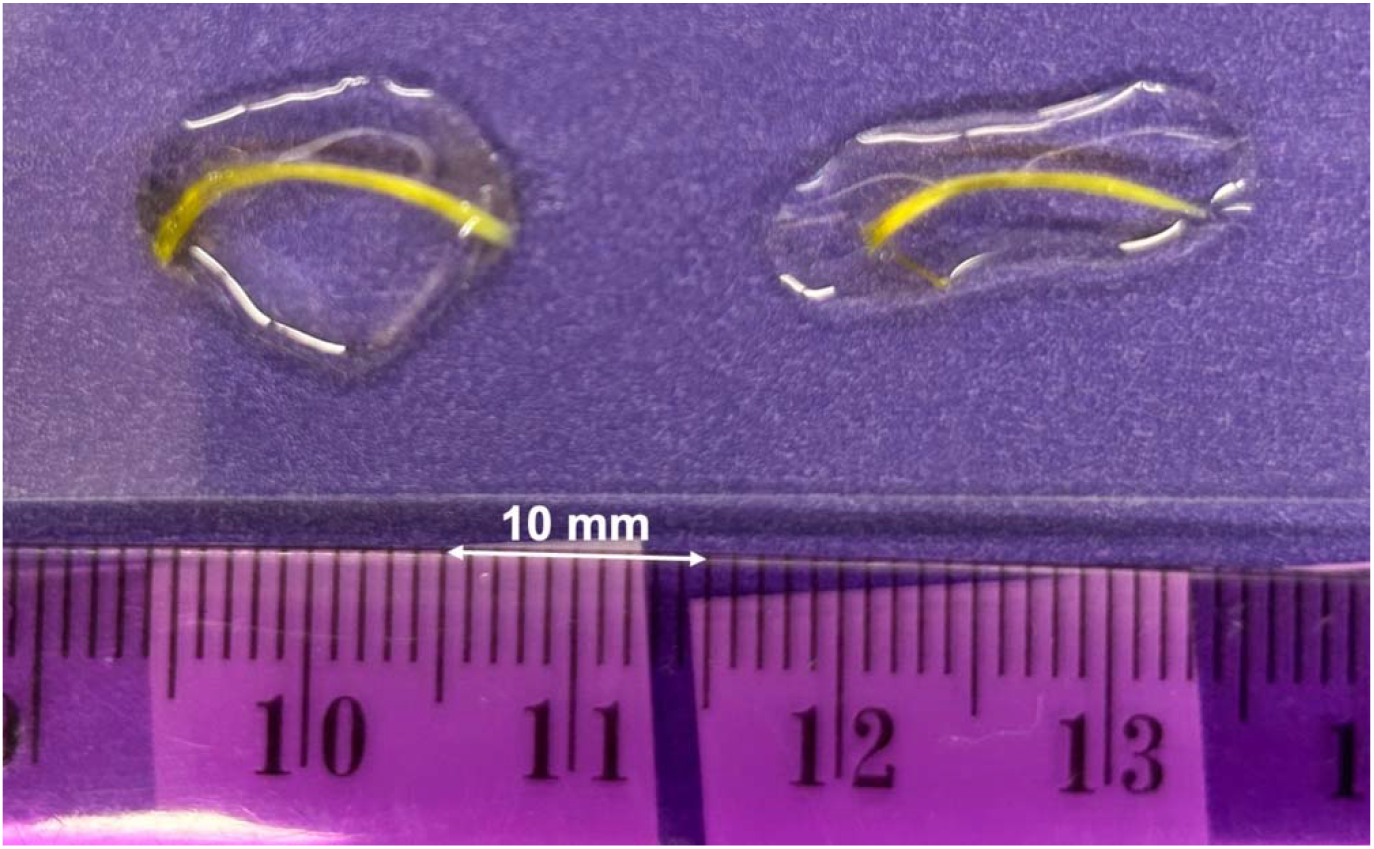
Example of distal fascicles from the right lateral gastrocnemius used for measurement of dissected fascicle length and calculation of serial sarcomere number, with fascicles positioned in the same plane as a ruler used to set the scale.

### Statistical Analysis

Statistical analyses were conducted using GraphPad Prism 9.5.1. To investigate variation in ultrasound-derived FL and PA, three-way analysis of variance (ANOVA) (time [pre-immobilization, post-immobilization] × joint position [90 degrees, full dorsiflexion] × region [proximal, middle, distal]) was performed for each muscle from each leg, with Geisser-Greenhouse corrections for sphericity. For each dissected muscle, a two-way ANOVA (leg [casted, un-casted] × region [proximal, middle distal] was used to investigate variation in SSN, SL, and FL, with Geisser-Greenhouse corrections for sphericity. For all ANOVAs, where interactions or effects of region were detected, pairwise comparisons (two-tailed paired t-tests) were performed with a Bonferonni correction for multiplicity. Two-tailed, paired t-tests compared muscle wet weights between the casted and un-casted leg, with a Bonferroni correction for multiplicity. For all significant t-tests, the effect size was reported as Cohen’s d. Significance was set at α = 0.05.

Linear regression was used to investigate the relationship between: 1) ultrasound-derived FL at 90° post-cast and FL of dissected fascicles; 2) ultrasound-derived FL at each joint angle post-cast and SSN of dissected fascicles; and 3) adaptations in ultrasound-derived FL (as % change pre to post-cast) at each joint angle and adaptations in SSN of dissected fascicles (as % change from the un-casted to the casted leg).

## Results

### Effects of region, joint position, and time on fascicle length measured via ultrasound

Three-way ANOVA results for FL measured via ultrasound are presented in Table 2.

**Table 2:** Three-way ANOVA results for ultrasound-derived fascicle length.

|  |  | Effect of region |  | Effect of joint position |  | Effect of time |  | Region × joint position interaction |  | Region × time interaction |  | Joint position × time interaction |  | Region × joint position × time interaction |  |
| --- | --- | --- | --- | --- | --- | --- | --- | --- | --- | --- | --- | --- | --- | --- | --- |
|  |  | F | P | F | P | F | P | F | P | F | P | F | P | F | P |
| LG | Un-casted | 393.80 | <0.0001* | 215.10 | <0.0001* | 3.99 | 0.0819 | 3.01 | 0.0655 | 0.08 | 0.921 | 17.79 | **0.0014\*** | 0.01 | 0.970 |
|  | Casted | 271.00 | <0.0001* | 269.10 | <0.0001* | 50.10 | <0.0001* | 2.43 | 0.106 | 3.02 | 0.0649 | 0.60 | 0.414 | 3.08 | 0.0954 |
| Soleus | Un-casted | 10.49 | **0.0022\*** | 148.90 | <0.0001* | 0.09 | 0.677 | 3.39 | **0.0480\*** | 2.07 | 0.145 | 1.13 | 0.296 | 0.67 | 0.426 |
|  | Casted | 24.83 | <0.0001* | 236.80 | <0.0001* | 27.64 | **0.0003\*** | 0.15 | 0.861 | 10.57 | **0.0004\*** | 0.17 | 0.662 | 0.26 | 0.629 |
| MG | Un-casted | 101.20 | <0.0001* | 99.57 | <0.0001* | 1.43 | 0.240 | 1.39 | 0.266 | 8.75 | **0.0011\*** | 2.70 | 0.132 | 1.69 | 0.214 |
|  | Casted | 120.30 | <0.0001* | 159.80 | <0.0001* | 3.62 | 0.0844 | 1.03 | 0.369 | 6.91 | **0.0036\*** | 14.30 | **0.0046\*** | 4.13 | 0.0596 |

For all muscles, there were effects of joint position, with FL increasing from a 90° ankle angle to full dorsiflexion (Table 2; Figures 2-4). For the gastrocnemii, there were effects of region, with FL increasing from proximal to distal (Table 2; Figures 2 and 4).

**Figure 2:**
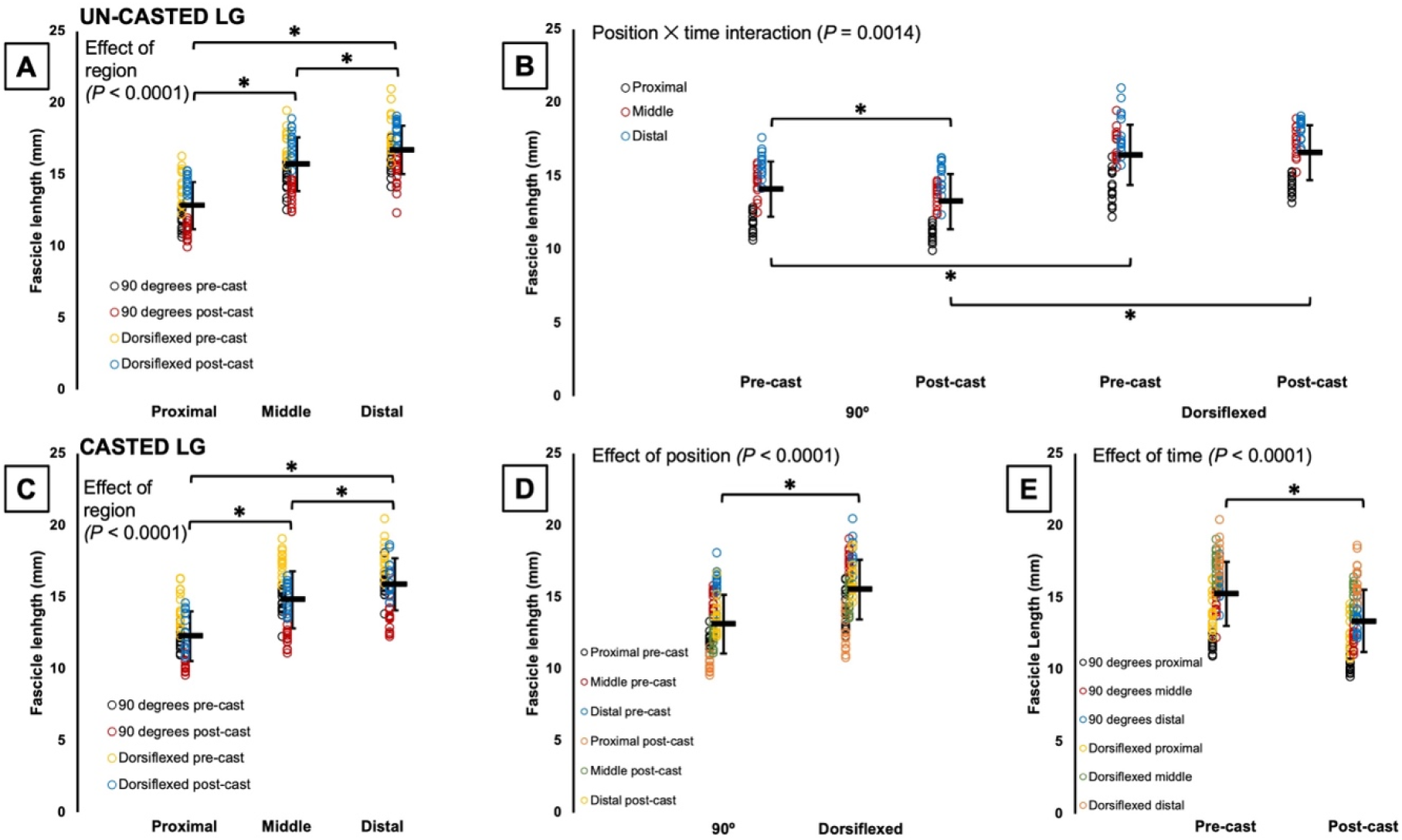
Fascicle length of the un-casted and casted lateral gastrocnemius (LG) measured via ultrasound. For the un-casted LG, there was an effect of region (A) and an interaction between joint position and time (B). For the casted LG, there were effects of region (C), joint position (D), and time (E). *Significant difference between indicated means (*P* < 0.05). Data are presented as mean ± standard deviation.

For ultrasound-derived FL of the un-casted LG, there was a joint position × time interaction (Table 2). Pairwise comparisons showed that FL decreased by 6% pre to post-cast when measurements were performed at 90° (P = 0.0001, *d* = 0.45), but did not change according to measurements performed at full dorsiflexion (P = 1.00) (Figure 2B).

For ultrasound-derived FL of the casted LG, there was an effect of time (Table 2), with FL decreasing by 12% pre to post-cast (Figure 2C).

For ultrasound-derived FL of the un-casted soleus, there was a region × joint position interaction (Table 2). Pairwise comparisons showed distal fascicles were longer than proximal fascicles when measured at 90° (P = 0.0413, *d* = 0.44), but proximal and middle FL did not differ (P = 0.194), and middle and distal FL did not differ (P = 1.00) (Figure 3A). Conversely, in measurements performed at full dorsiflexion, middle fascicles were longer than proximal fascicles (P = 0.0003, *d* = 0.65), but proximal and distal FL did not differ (P = 1.00), and middle and distal FL did not differ (P = 0.0591) (Figure 3A). FL of the un-casted soleus did not change from pre to post-cast, with no effect of time (Figure 3B).

**Figure 3:**
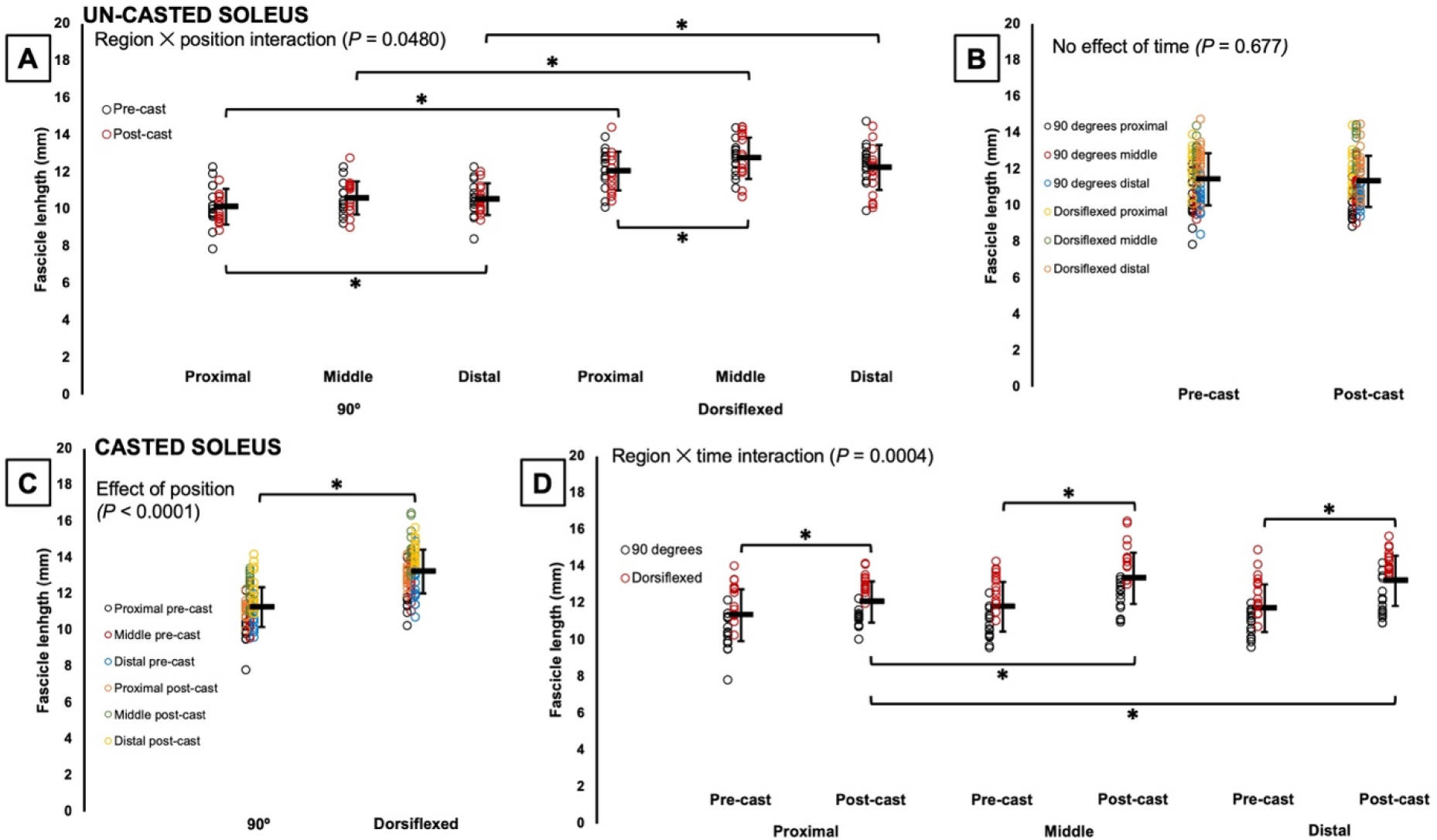
Fascicle length of the un-casted and casted soleus measured via ultrasound. For the un-casted soleus, there was an interaction between region and position (A) and no effect of time (B). For the casted soleus, there was an effect of position (C) and an interaction between region and time (D). *Significant difference between indicated means (*P* < 0.05). Data are presented as mean ± standard deviation.

For the casted soleus, an effect of time showed that ultrasound-derived FL increased on average by 11% pre to post-cast (Table 2). There was also a region × time interaction. Pairwise comparisons showed all regions of the soleus increased FL from pre to post-cast (proximal: P = 0.0301, *d* = 0.57; middle: P < 0.0001, *d* = 1.12; distal: P < 0.0001, *d* = 1.13) (Figure 3D). Pre-cast, there were no regional differences in FL (P = 0.0849-1.00), but post-cast, middle (P < 0.0001; *d* = 1.01) and distal fascicles (P < 0.0001; *d* = 0.91) were longer than proximal fascicles (Figure 3D). Accordingly, the increase in proximal FL from pre to post-cast was smaller (+6%) than the increases in middle and distal FL (both +13%).

For ultrasound-derived FL of the un-casted MG, there was a region × time interaction (Table 2), with distal FL increasing by 8% pre to post-cast (P = 0.0330, *d* = 0.63) (Figure 4A).

**Figure 4:**
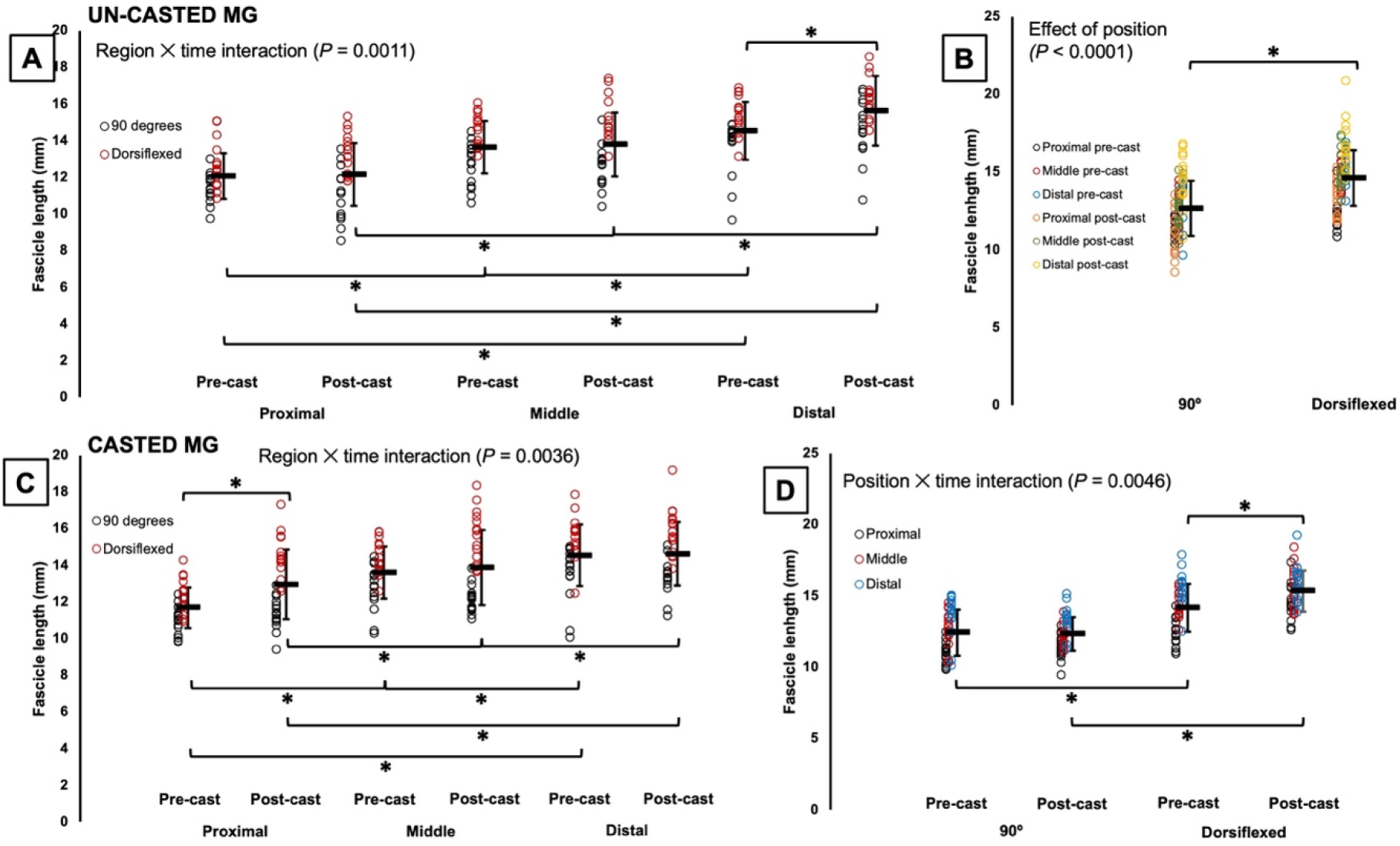
Fascicle length of the un-casted and casted medial gastrocnemius (MG) measured via ultrasound. For the un-casted MG, there was an interaction between region and time (A) and an effect of position (B). For the casted MG, there were interactions between region and time (C) and position and time (D). *Significant difference between indicated means (*P* < 0.05). Data are presented as mean ± standard deviation.

For ultrasound-derived FL of the casted MG, there was also a region × time interaction (Table 2), but with proximal FL increasing by 11% pre to post-cast (P = 0.0028, *d* = 0.82) (Figure 4C). A joint position × time interaction showed that measurements at 90° detected no change in FL pre to post-cast (P = 1.00), but measurements at full dorsiflexion detected an 8% increase in FL (P = 0.0002, *d* = 0.76) (Figure 4D).

### Effect of time on pennation angle measured via ultrasound

Three-way ANOVA results for PA measured via ultrasound are presented in Table 3. For the un-casted LG, there was an effect of time (Table 3) such that PA increased by ∼10% pre to post-cast (Figure 5A). For the casted LG, there was a region × joint position × time interaction (Table 3). Pairwise comparisons showed a 26% decrease in PA pre to post-cast only in distal fascicles at full dorsiflexion (P < 0.0001, *d* = 3.24) (Figure 5B).

**Figure 5:**
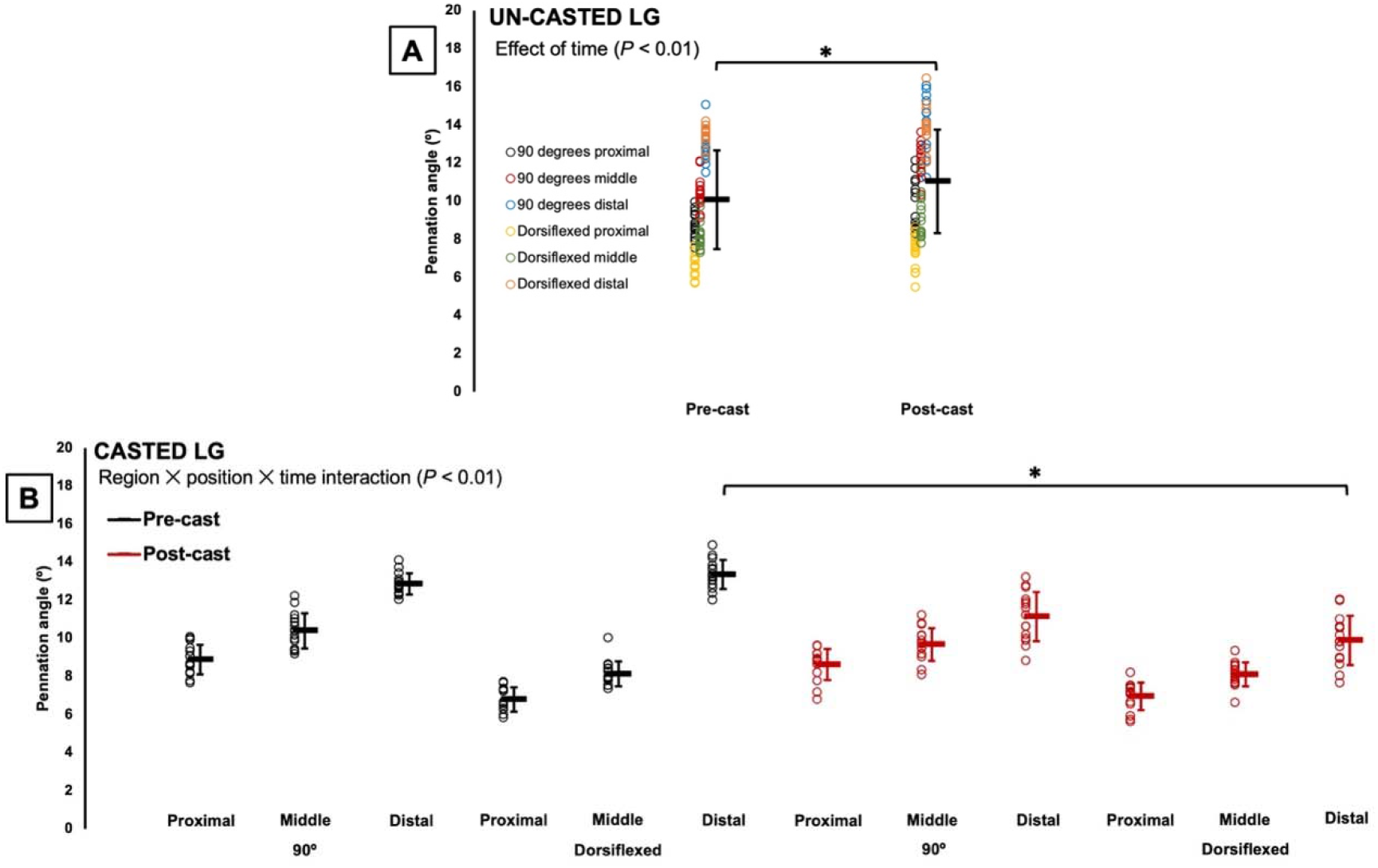
Changes in pennation angle of the un-casted (A) and casted (B) lateral gastrocnemius (LG) from pre to post-cast. *Significant difference between indicated means (*P* < 0.05). Data are presented as mean ± standard deviation.

**Table 3:**
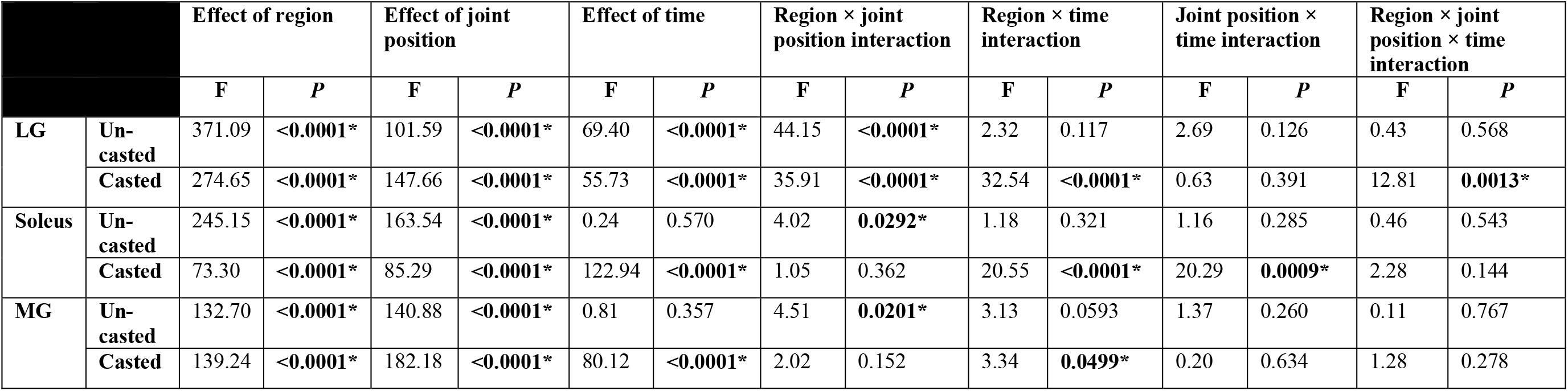
Three-way ANOVA results for ultrasound-derived pennation angle

For the un-casted soleus, like with FL, time did not affect PA, with no changes pre to post-cast (Table 3; Figure 6A). For the casted soleus, there were interactions of region × time and joint position × time (Table 3). Pairwise comparisons showed that at all regions of muscle, and both joint angles, PA of the casted soleus decreased (9-31%) pre to post-cast (P < 0.0001-0.0005, *d* = 1.08-2.78) (Figure 6B-C).

**Figure 6:**
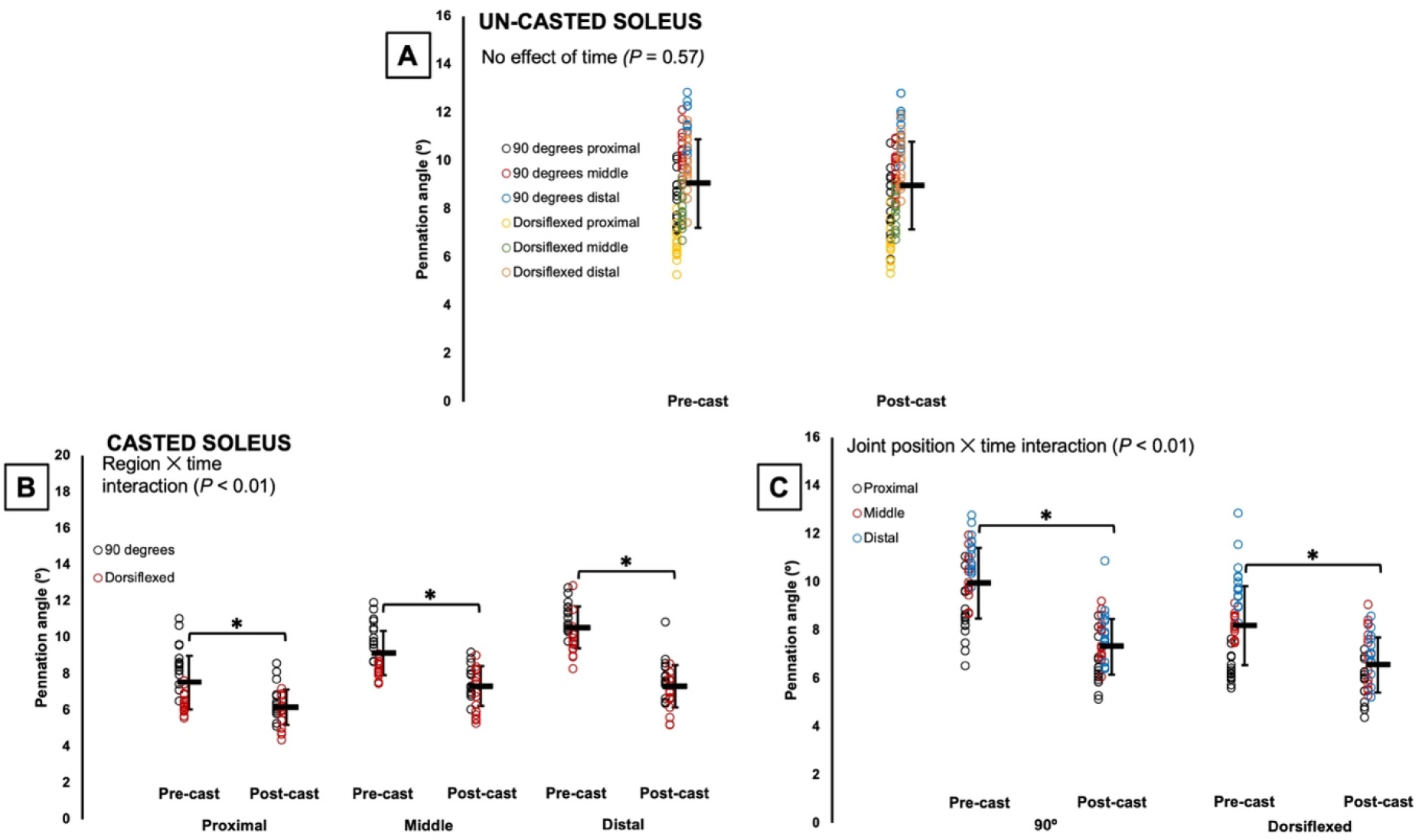
Changes in pennation angle of the un-casted (A) and casted (B-C) soleus from pre to post-cast. *Significant difference between indicated means (*P* < 0.05). Data are presented as mean ± standard deviation.

For the un-casted MG, time did not affect PA, with no changes pre to post-cast (Table 3; Figure 7A). For the casted MG, there was a region × time interaction (Table 3), and pairwise comparisons showed that at all three regions of the muscle, PA decreased by ∼20% pre to post-cast (P < 0.0001-0.0003, *d* = 1.10-2.05) (Figure 7B).

**Figure 7:**
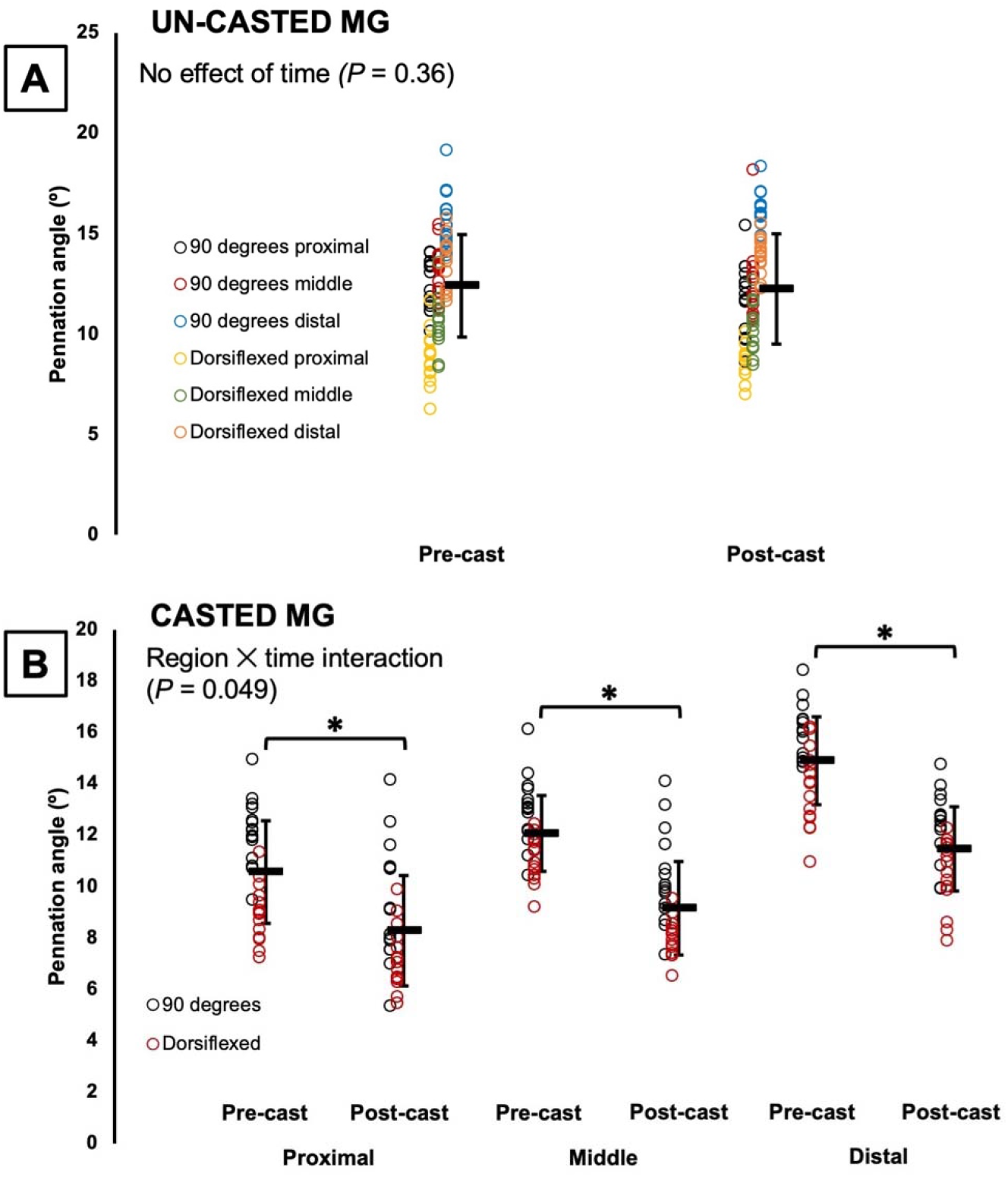
Changes in pennation angle of the un-casted (A) and casted (B) medial gastrocnemius (MG) from pre to post-cast. *Significant difference between indicated means (*P* < 0.05). Data are presented as mean ± standard deviation.

### Muscle wet weight in the casted versus un-casted leg

The LG, soleus, and MG of the casted leg weighed 62%, 33%, and 54% less, respectively, than the muscles of the un-casted leg (P < 0.0001, *d* = 2.42-2.96) (Figure 8).

**Figure 8:**
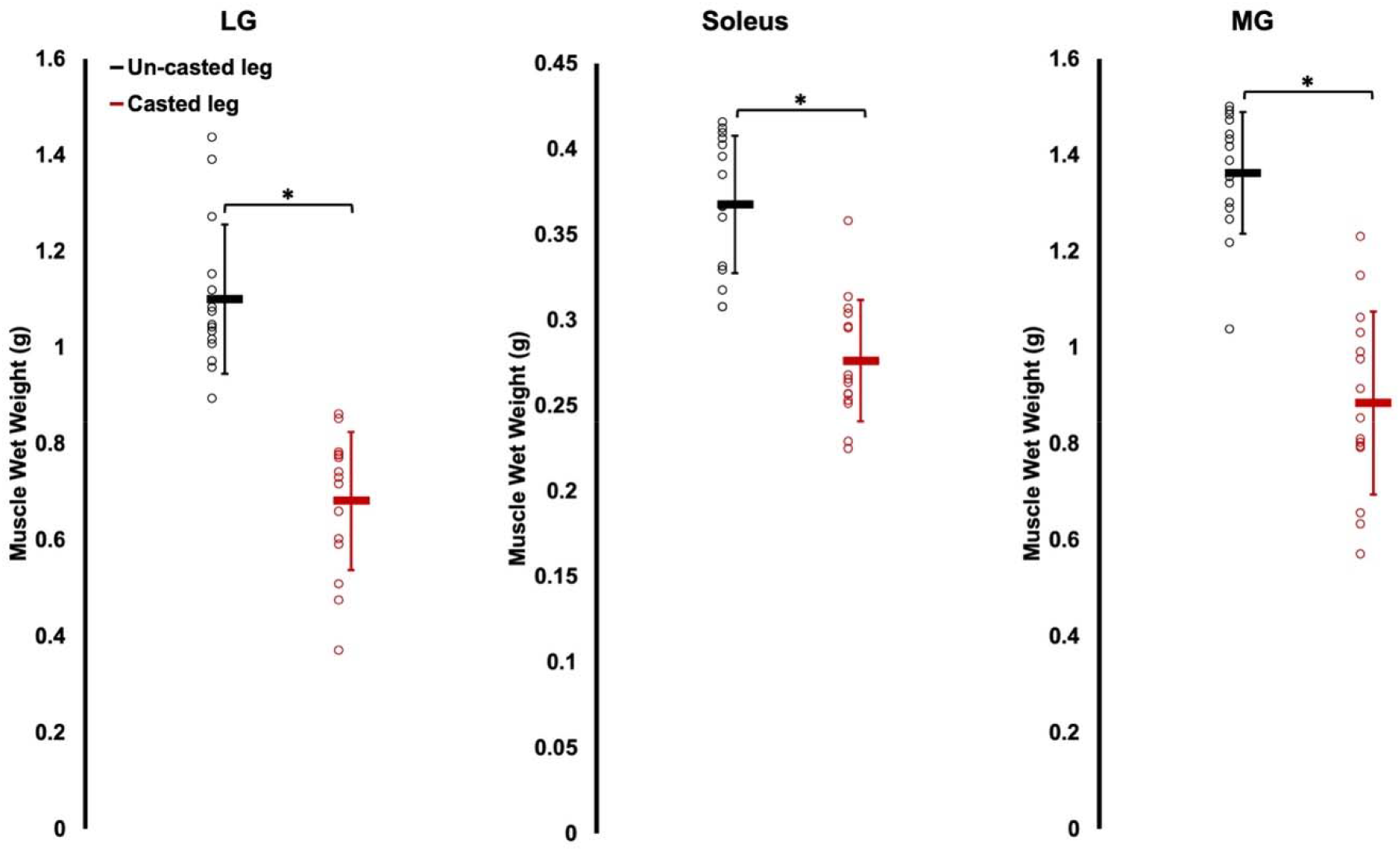
Comparison of muscle wet weight between the casted and un-casted leg for the lateral gastrocnemius (LG), soleus, and medial gastrocnemius (MG). *Significant difference between indicated means (*P* < 0.05). Data are presented as mean ± standard deviation.

### Serial sarcomere number, sarcomere length, and fascicle length of the dissected fascicles in the casted versus non-casted leg

Two-way ANOVA results for SSN, SL, and FL of the dissected fascicles are shown in Table 4. There were no region × leg interactions for any muscles.

**Table 4:** Two-way ANOVA results for serial sarcomere number, sarcomere length, and fascicle length of dissected fascicles.

|  |  | Effect of region |  | Effect of leg |  | Region × leg interaction |  |
| --- | --- | --- | --- | --- | --- | --- | --- |
|  |  | F | P | F | P | F | P |
| LG | SSN | 108.91 | <0.0001* | 4.57 | 0.0507 | 0.67 | 0.502 |
|  | SL | 61.46 | <0.0001* | 8.37 | 0.0118* | 2.77 | 0.0959 |
|  | FL | 55.00 | <0.0001* | 11.57 | 0.0043* | 0.17 | 0.793 |
| Soleus | SSN | 1.39 | 0.267 | 33.02 | <0.0001* | 0.68 | 0.475 |
|  | SL | 1.26 | 0.291 | 0.14 | 0.713 | 0.64 | 0.514 |
|  | FL | 3.06 | 0.0639 | 22.94 | 0.0003* | 0.10 | 0.871 |
| MG | SSN | 21.30 | <0.0001* | 0.99 | 0.336 | 1.01 | 0.374 |
|  | SL | 18.36 | <0.0001* | 6.27 | 0.0253* | 1.02 | 0.373 |
|  | FL | 7.77 | 0.0111* | 5.03 | 0.0416* | 0.62 | 0.541 |

For the LG, there were effects of leg (Table 4) on SL and FL of dissected fascicles such that they were 3% and 6% shorter, respectively, in the casted LG (Figure 9B-C). SSN did not differ between the casted and un-casted LG (Figure 9A). There were effects of region on SSN, SL, and FL. SSN increased from proximal to middle to distal (P < 0.0001, *d* = 1.30-3.24) (Figure 9D), and FL followed a similar trend (P < 0.0001, *d* = 1.58-2.04) but with no difference between proximal and middle FL (P = 0.0521) (Figure 9F). Conversely, SL decreased from proximal to middle to distal (P < 0.0001-0.0016, *d* = 0.90-2.41) (Figure 9E).

**Figure 9:**
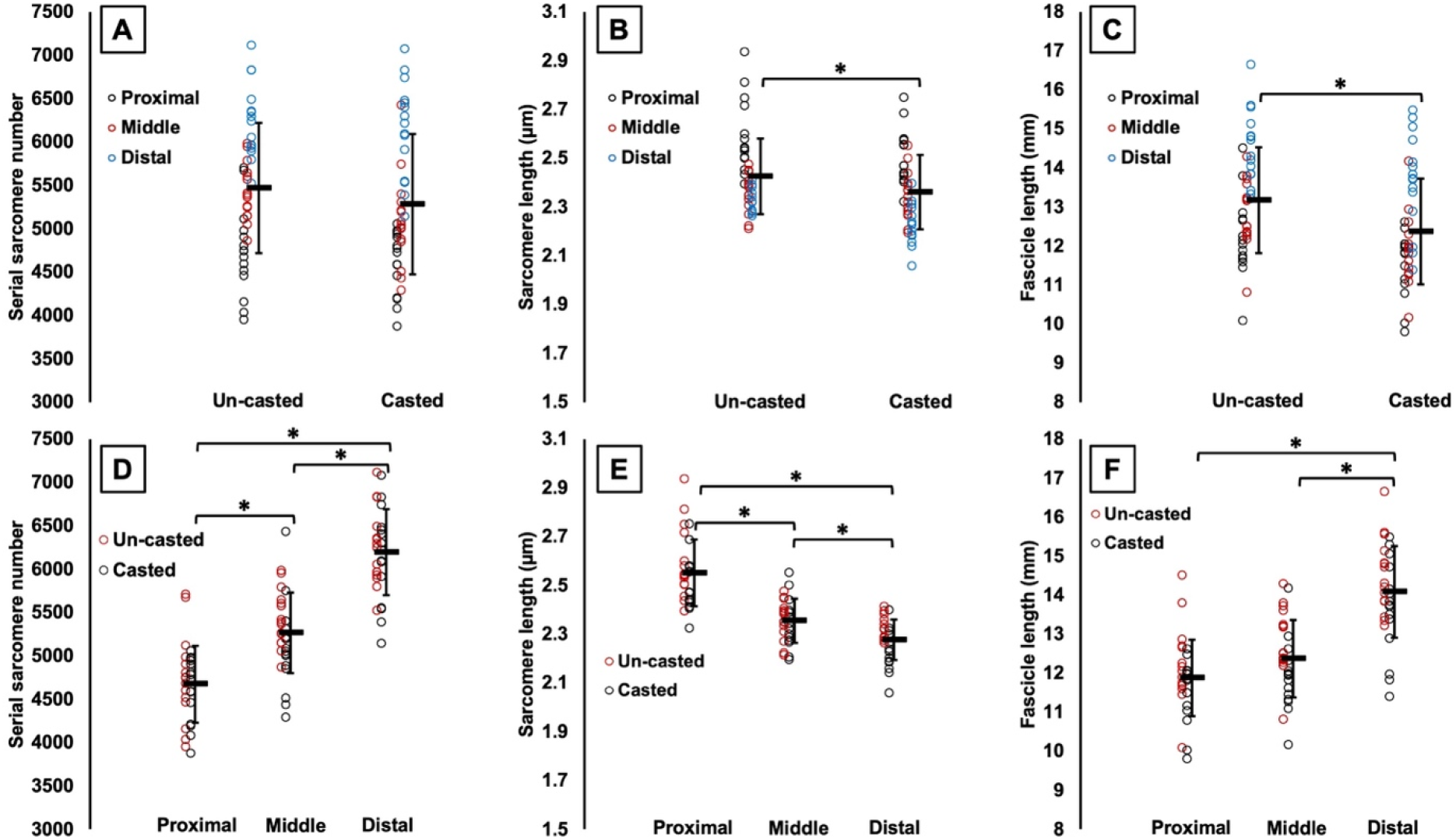
Effects of time (A to C) and effects of region (D to F) on serial sarcomere number (A and D), sarcomere length (B and E), and fascicle length (C and F) of the lateral gastrocnemius from dissected fascicles. *Significant difference between indicated means (*P* < 0.05). Data are presented as mean ± standard deviation.

For the soleus, there were no effects of region on SSN, SL, or dissected FL, indicating no regional differences (Table 4). There was an effect of leg on soleus SSN (Table 4) such that SSN was 6% greater in the casted leg (Figure 10A). There was a similar effect of leg on FL (Table 4), with a 6% increase (Figure 10C). Soleus SL did not differ between the casted and un-casted leg (Table 4; Figure 10B).

**Figure 10:**
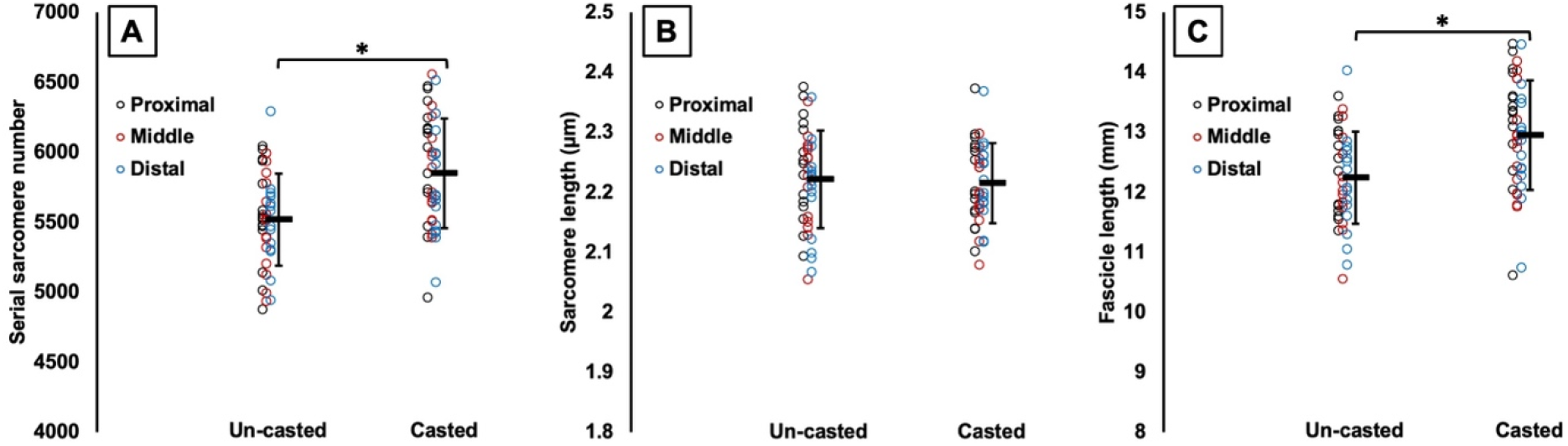
Effects of time on serial sarcomere number (A), sarcomere length (B), and fascicle length (C) of the soleus from dissected fascicles. *Significant difference between indicated means (*P* < 0.05). Data are presented as mean ± standard deviation.

For the MG, there were effects of leg on SL and dissected FL (Table 4) such that they were 2% and 4% shorter, respectively, in the casted MG (Figure 11B-C). SSN did not differ between the casted and un-casted MG (Figure 11A). There were effects of region on SSN, SL, and FL (Table 4). SSN followed the same pattern as in the LG, increasing from proximal to middle to distal (P < 0.0001-0.0127, *d* = 0.79-1.42) (Figure 11D). FL only differed between proximal and distal fascicles, with distal fascicles being longer (P = 0.0044, *d* = 0.43) (Figure 11F). Like with the LG, SL decreased from proximal to middle to distal (P < 0.0001-0.0481, *d* = 0.52-1.49) (Figure 11E).

**Figure 11:**
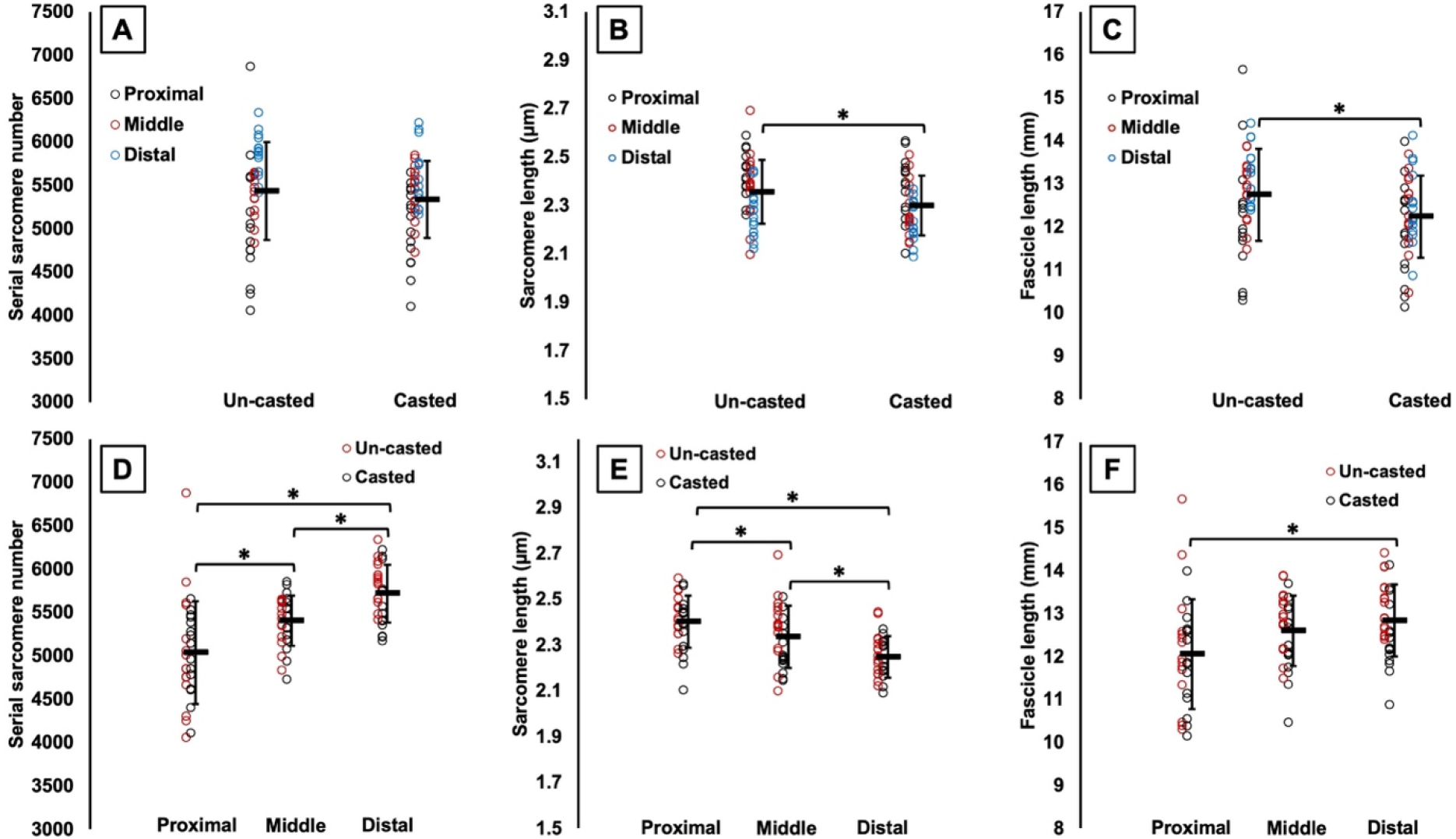
Effects of time (A to C) and effects of region (D to F) on serial sarcomere number (A and D), sarcomere length (B and E), and fascicle length (C and F) of the medial gastrocnemius from dissected fascicles. *Significant difference between indicated means (*P* < 0.05). Data are presented as mean ± standard deviation.

### Relationships between adaptations in fascicle length measured via ultrasound and adaptations in serial sarcomere number and fascicle length measured from dissected fascicles

For the soleus, significant positive relationships were found between ultrasound-derived FL at 90° and FL of dissected fascicles (Figure 12A) and SSN (Figure 12D), and between ultrasound-derived FL at full dorsiflexion and SSN (Figure 12G). For the LG, there was only a relationship between ultrasound-derived FL at 90° and FL of dissected fascicles (Figure 12B), and no relationships among these measures were observed for the MG (Figure 12C, F, I).

**Figure 12:**
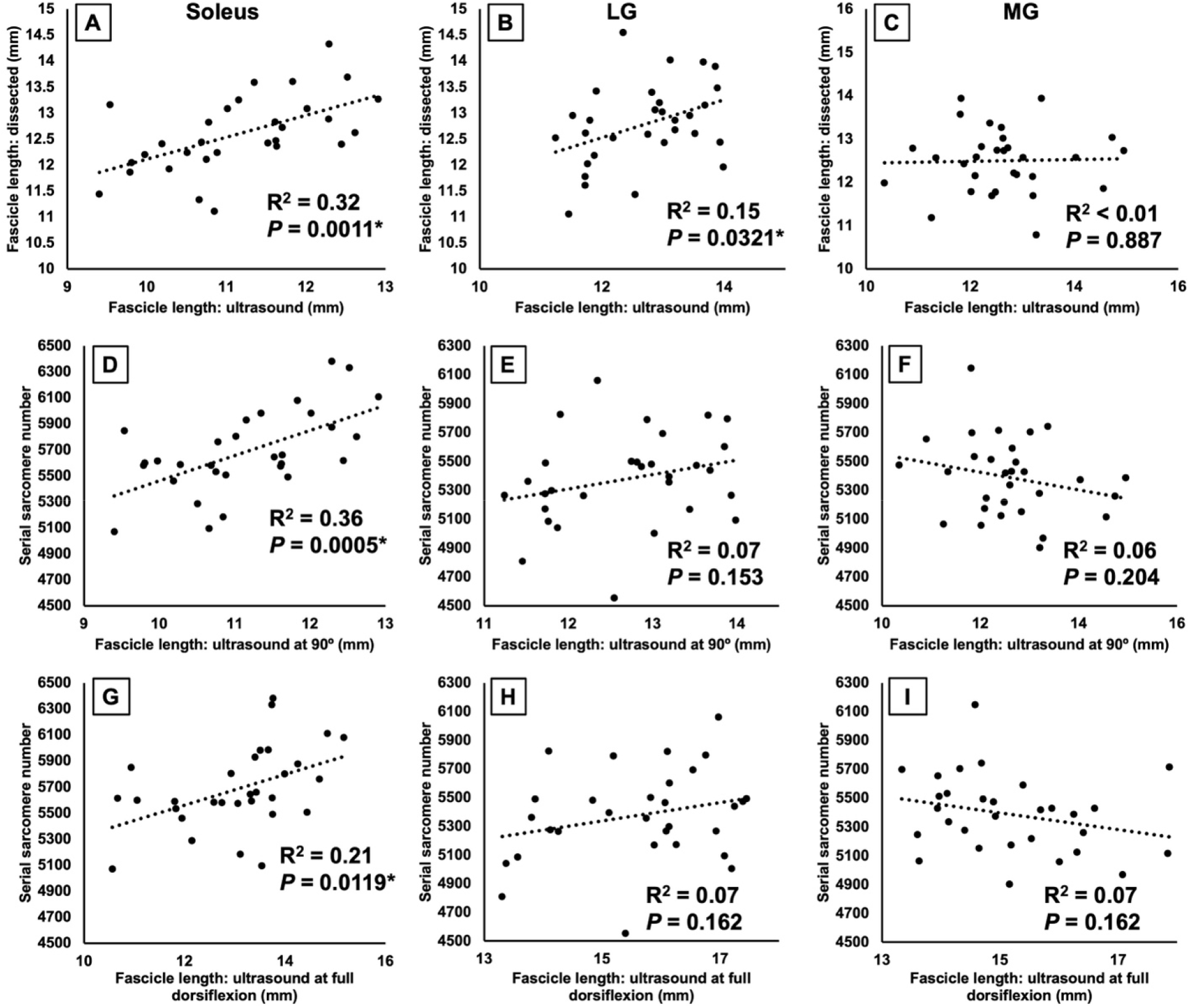
Relationships between ultrasound-derived fascicle length at 90° and fascicle length of dissected fascicles (A-C), ultrasound-derived fascicle length at 90° and SSN (D-F), and ultrasound-derived FL at full dorsiflexion and SSN (G-I) for the soleus, lateral gastrocnemius (LG), medial gastrocnemius (MG). *Significant relationship (P < 0.05).

There were no relationships between the % change in ultrasound-derived FL from pre to post-cast and the % change in SSN from the un-casted to casted leg for the LG, soleus, or MG (Table 5).

**Table 5:**
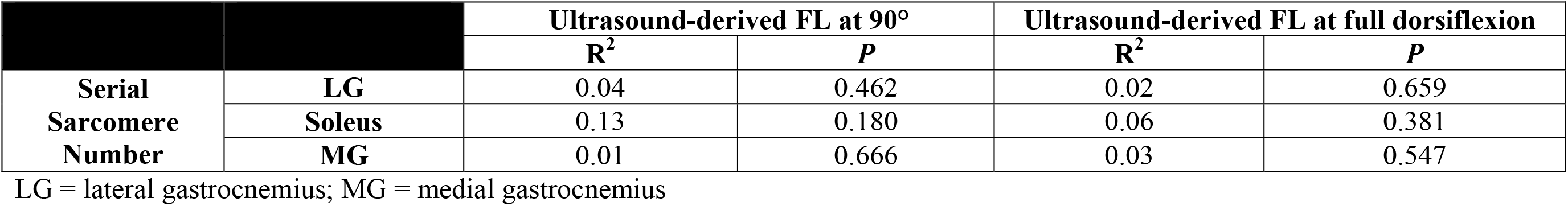
Relationships between % change in ultrasound-derived fascicle length (FL) from pre to post-cast and % change in serial sarcomere number of dissected fascicles from the un-casted to casted leg

When regression analyses were performed across all muscles together, the % change in ultrasound-derived FL measured with the ankle at 90° explained 28% of the variation in the % change in SSN from dissected fascicles (Figure 13A). This relationship was lessened when using ultrasound-derived FL at full dorsiflexion, only explaining 10% of the variation in SSN adaptations (Figure 13B).

**Figure 13:**
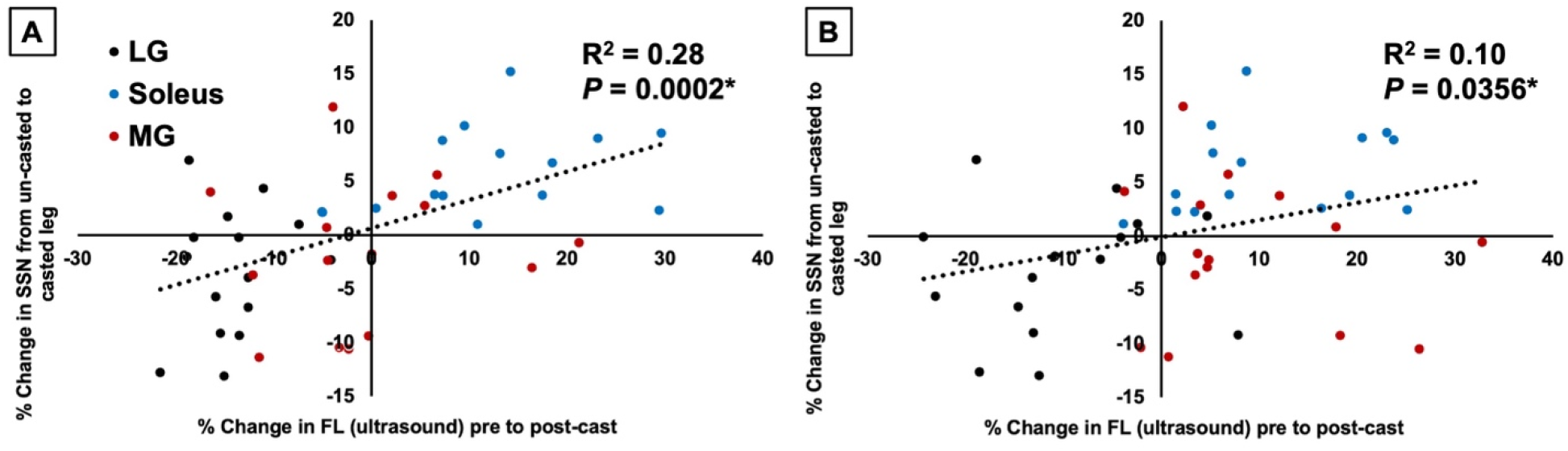
For all muscles combined, relationships between the % change in ultrasound-derived fascicle length (FL) from pre to post-cast as measured with the ankle at 90° (A) and full dorsiflexion (B) and the % change in serial sarcomere number (SSN) from the un-casted to casted leg determined from dissected fascicles. *Significant relationship (P < 0.05).

## Discussion

Immobilizing the rat ankle in full dorsiflexion for 2 weeks, the present study investigated whether ultrasound-derived FL measurements can accurately depict SSN adaptations. Ultrasound detected an 11% increase in soleus FL, a 12% decrease in LG FL, and (depending on the joint angle and region of muscle) an 8% increase in MG FL. These adaptations were partly reflected by the SSN measurements obtained from dissected fascicles, with a 6% greater soleus SSN in the casted leg than the un-casted leg, but no differences in SSN for the gastrocnemii. Our results indicate that ultrasonographic measurements of FL can overestimate SSN adaptations.

Our values for muscle wet weight, SSN, SL, FL, and ultrasound-derived PA are within previously reported ranges for the rat soleus (Booth, 1977; Soares *et al*., 2007; Peixinho *et al*., 2011; Mele *et al*., 2016; Chen *et al*., 2020; Hinks *et al*., 2022*b*) and gastrocnemii (Booth, 1977; Woittiez *et al*., 1986; Ochi *et al*., 2007; Peixinho *et al*., 2011; Mele *et al*., 2016; Power *et al*., 2021).

### Stretch-induced adaptation in serial sarcomere number

The soleus of the casted leg had a 6% greater SSN than the un-casted leg, which is consistent with findings from previous studies that immobilized the soleus in a stretched position in rats, mice, and cats (Tabary *et al*., 1972; Williams & Goldspink, 1978; Spector *et al*., 1982; Shah *et al*., 2001; Soares *et al*., 2007; Kinney *et al*., 2017). This serial sarcomere addition is believed to occur to restore optimal actin-myosin overlap and reduce sarcomeric passive tension in the stretched position (Williams & Goldspink, 1978; Davis *et al*., 2020; Hinks *et al*., 2022*a*). The increase in rat soleus SSN we observed after 2 weeks of immobilization was notably lower (+6%, from 5518 to 5850 sarcomeres) than the increase reported by Soares *et al*. (2007) after immobilizing the soleus in a stretched position for just 4 days (+29%, 6338 to 8174 sarcomeres). This discrepancy may be attributed to the more extreme ankle angle they used for immobilization, described as “total dorsiflexion,” while we immobilized the ankle at 40°. Additionally, we did not observe a difference in SSN between the casted and un-casted legs for the gastrocnemii. The gastrocnemii are biarticular muscles, crossing the ankle and the knee. Spector *et al*. (1982) immobilized the knee in full extension and the ankle at 45°, and observed a 20% increase in MG SSN. While we immobilized the ankle at a similar angle (40°), we did not immobilize the knee, allowing movement of the gastrocnemii at that joint, which likely tempered the stretch stimulus imposed by dorsiflexion for those muscles.

### Immobilization-induced atrophy

Our casting intervention induced atrophy in the gastrocnemii and soleus, as evidenced by lower muscle wet weights in the casted leg. Measurements of PA provided by ultrasound align with these findings, showing decreased PA from pre to post-cast in all three muscles, which may reflect the loss of sarcomeres in parallel (Wisdom *et al*., 2015; Jorgenson *et al*., 2020). The reduced muscle weight was more pronounced in the gastrocnemii (–54-62%) than the soleus (– 33%). Considering the soleus had a 6% greater SSN in the casted than the un-casted leg, an increase in SSN due to stretch may have lessened the overall loss of muscle tissue, limiting the loss to only sarcomeres in parallel. A similar result was observed by Spector *et al*. (1982), with a smaller reduction in soleus wet weight when immobilizing in a stretched position (–14%) than a shortened position (–48%), and the former increasing SSN while the latter decreased SSN. The gastrocnemii in the present study did not appear to experience an increase in SSN, therefore, the loss of parallel sarcomeres may not have been made up for by stretch-induced serial sarcomere addition, resulting in a greater loss of muscle weight.

### Can the un-casted leg be used as a valid control?

In the un-casted soleus, no differences in FL or PA were detected by ultrasound from pre to post-cast, validating the use of the un-casted soleus as a SSN control in the present study and previous studies (Williams & Goldspink, 1978; Heslinga & Huijing, 1993; Shah *et al*., 2001; Gomes *et al*., 2004; Kinney *et al*., 2017). In the un-casted gastrocnemii, however, ultrasound measurements suggest some adaptations may have occurred, possibly due to the un-casted leg compensating for the added load (the cast) on the opposite leg during ambulation. In the un-casted LG, ultrasound showed a 6% decrease in FL at 90°, accompanied by a 10% increase in PA. Increased PA and sometimes a decrease in FL are often observed following training emphasizing concentric contractions, and may reflect a reorganization of the muscle architecture to add sarcomeres in parallel for greater force production (Butterfield *et al*., 2005; Franchi *et al*., 2014).

### Ultrasound-derived FL does not perfectly reflect adaptations in serial sarcomere number

Measurements of FL via ultrasound are often used to infer increases or decreases in SSN (Narici *et al*., 2003; Blazevich *et al*., 2007; Franchi *et al*., 2014; Hinks *et al*., 2021). Inferring SSN adaptations from ultrasound-derived FL may be problematic, however, because apparent changes in FL may simply be due to changes in SL at the joint angle at which ultrasound measurements are obtained. For example, Pincheira *et al*. (2021) observed an increase in biceps femoris FL following 3 weeks of eccentric training as measured with the leg in full extension; however, microendoscopy revealed the increase in FL was only due to longer SLs at that joint angle, not training-induced serial sarcomere addition. In research on animals, SSN adaptations are often determined by calculating SSN from measurements of SL and FL from dissected fascicles, then comparing between experimental and control muscles. The present study investigated the relationship between these two most commonly used methodologies for assessing SSN adaptations.

We observed significant but weak relationships between ultrasound-derived FL at 90° and FL of dissected fascicles (after being fixed at 90°) for the soleus and LG (Figure 12A-B). *Kellis et al.* (2009) observed moderate to strong relationships between FL measured via ultrasound and FL measured directly in the hamstrings of human cadavers. Our results may differ from theirs because, after digesting the muscles in nitric acid, it was more difficult to ensure that the same fascicles as the ultrasound images were being measured from the dissected muscle, even though the same regional constraints (two fascicles from each of the proximal, middle, and distal regions) were followed. Additionally, we observed relationships between ultrasound-derived measurements of FL and actual SSN determined from dissected fascicles for the soleus only, and between the % change in FL from pre to post-cast and the % change in SSN from the un-casted to casted leg with all muscles together. In both cases, the relationships using FL measured at full dorsiflexion were weaker than when using FL measured at 90°. Similar findings were observed recently by Werkhausen *et al*. (2023), with the relationship between ultrasound-derived FL and isokinetic force (i.e., a measure associated with SSN (Drazan *et al*., 2019; Hinks *et al*., 2022*a*)) being moderate or non-existent depending on the joint angle used during ultrasound imaging. Collectively, our regression analyses demonstrate variability both among muscles and between joint angles in the ability for ultrasound-derived FL to truly represent SSN.

Overall, the soleus provided the best means for comparing ultrasound-derived FL adaptations and adaptations in SSN, as the un-casted soleus did not appear to undergo any compensatory adaptations. From the ultrasound measurements, we observed an ∼11% increase in soleus FL from pre to post-cast, however, the true increase in SSN from the un-casted to the casted leg was only 6%. This serial sarcomere addition appeared to be driven by a 6% increase in FL, as the un-casted and casted soleus had the same SL (∼2.2 µm) with the ankle fixed at 90°. Interestingly, while ultrasound-derived FL averaged across muscle regions increased by 11%, the increase in ultrasound-derived FL of proximal fascicles (+6%) was closer to the observed increase in SSN, demonstrating regional variability in the accuracy of ultrasound-derived FL measurements. Altogether, an increase in FL measured by ultrasound can indeed correspond to an increase in SSN in the rat soleus, but may overestimate the increase in SSN by as much as 5%.

### Limitations of ultrasound that may contribute to a disconnect between ultrasound-derived FL and actual SSN

For the gastrocnemii, the distal fascicles were sometimes partly out of plane (Figure 1), thus the trajectory of those fascicles to the deep aponeurosis was used to complete the measurements of FL. This limitation likely contributed to the higher coefficients of variation for the gastrocnemii compared to the soleus (Table 1), and may explain the lack of relationships observed between ultrasound-derived FL and dissected FL and SSN for the gastrocnemii, but not the soleus. It is also important to note that ultrasound images do not capture the contractile tissue of muscle fascicles, but rather the perimysium, the sheath of connective tissue surrounding each fascicle. During serial sarcomerogenesis, the connective tissue scaffolding must be constructed before sarcomeres are added within that space (Kjær, 2004). Previous studies have reported no changes (Williams *et al*., 1988) or increases in intramuscular connective tissue content (Ahtikoski *et al*., 2001) following immobilization in a stretched position depending on the duration of immobilization. Many studies also overlook that adaptations in connective tissue structure (e.g., crosslinking, collagen fibril orientation, organization) may not be evident in measures of only content, but can change during muscle remodelling as well (Kjær, 2004). Connective tissue is digested in nitric acid before measurements are performed on dissected fascicles, therefore, variability in connective tissue likely affects the ability for ultrasound to capture FL of only contractile tissue. Furthermore, an ultrasound image only captures a fascicle path in two dimensions, but the three-dimensional nature of fascicle curvature is well-documented (Rana *et al*., 2013; Raiteri *et al*., 2016; Cameron *et al*., 2023). Unless methods such as three-dimensional ultrasound (Raiteri *et al*., 2016) or magnetic resonance diffusion tensor imaging (Cameron *et al*., 2023) are used, the three-dimensional nature of FL can only be accounted for when fascicles are dissected out of the muscle. In the present study, this two-dimensional limitation of ultrasound is most evident in how dissected FLs of the soleus were ∼13% longer than ultrasound-derived FLs. There may be curvature in rat soleus fascicles that is not captured in a lateral ultrasound scan, making fascicles appear shorter. Altogether, these factors may have contributed to the disconnects between ultrasound-derived FL and actual SSN in the present study, including the ∼5% overestimation of sarcomerogenesis in the soleus, and should be considered going forward in studies employing muscle ultrasound.

## Conclusion

The present study investigated the relationship between the two most commonly used methods of assessing longitudinal growth of skeletal muscle: 1) ultrasound-derived FL measurements pre and post-intervention; and 2) comparison of SSN between an experimental and a control muscle. We showed that ultrasound-derived FL overestimated SSN adaptations by ∼5%, with measurements in a neutral position predicting SSN better than measurements in a stretched position. Future studies should consider these findings when concluding a large magnitude of serial sarcomerogenesis based on ultrasound-derived FL taken at a set joint angle, and may consider applying a correction factor to more closely approximate the actual SSN adaptations.

## Acknowledgements

This project was supported by the Natural Sciences and Engineering Research Council of Canada (NSERC). No conflicts of interest, financial or otherwise, are declared by the authors.

## Ethics statement

Approval was given by the University of Guelph’s Animal Care Committee and all protocols followed CCAC guidelines (AUP #4905).

## Data accessibility

Individual values of all supporting data are available upon request.

## Author contributions

A.H., M.V.F. and G.A.P. conceived and designed research; A.H. carried out animal husbandry and training; A.H. performed experiments; A.H. analyzed data; A.H., M.V.F., and G.A.P. interpreted results of experiments; A.H. prepared figures; A.H. and G.A.P. drafted manuscript; A.H., M.V.F., and G.A.P. edited and revised manuscript; A.H., M.V.F., and G.A.P. approved final version of manuscript.

